# The changes in blood flow velocity in the internal jugular vein under conditions of variable intrapulmonic pressure and gravitational effect

**DOI:** 10.1101/2023.07.28.551064

**Authors:** Anton Kasatkin, Aleksandr Urakov, Ivan Kuvshinov, Nikita Zhulkov

## Abstract

**Background:** We explored the dynamics of changes in the blood flow velocity in the internal jugular vein (IJV) under conditions of changing intrapulmonary pressure and gravitational influence in healthy volunteers.

**Methods:** Color duplex ultrasonography of the right IJV was performed in 25 healthy volunteers of both genders. The dopplergram of the blood flow spectrum was recorded in the right IJV in the horizontal supine position of the participants, and 3 minutes after the volunteers had attained the new position due to the lifting the bed back section at the 30°. We recorded the maximum (Max) and minimum (Min) linear of blood flow velocity (cm/s). In each position, the measurements were carried out sequentially at positive end-expiratory pressure (PEEP) values of 0, 5, 10 cmH2O after 3 minutes by process of changing the level of PEEP. Quantitative data were presented as arithmetic mean (M), standard deviation (SD) and compared using the Wilcoxon rank-sum test.

**Results:** Our data demonstrated that negatively related influence of continuous positive airway pressure on IJV blood flow velocity: Max/Min 57.40±13.10/20.68±7.80 cm/s (PEEP 0 cmH2O), 39.00±18.80/14.63±4.90 (PEEP 5 cmH2O), 30.83±7.30/13.25±2.90 (PEEP 10 cmH2O). The blood flow velocity values are increased after 30° head elevation in PEEP 5 and 10 cmH2O in comparison to 0° values.

**Conclusions:** The constant positive airway pressure 5-10 cmH2O in a person in supine position decreasing blood flow velocity in IJV. The 30° head elevation eliminates this effect.

## Introduction

In the description of separate clinical cases, it will be founded that reduction of the blood flow velocity in the central veins can increase the risk of thrombosis.^1-3^ It is well-known, that reduced blood flow accompanied by hypercoagulation and defect of the vessel wall compound go to make up, what is called Virchow’s triad, leads to thrombosis.^4,5^ Internal jugular vein (IJV) thrombus can represent a medical problem, determinative quality and safety of health care. In intensive care unit, an incidence of IJV thrombosis with clinical symptomatic can reach 2-5% and asymptomatic – 33-67%.^6^ The etiology of the thrombosis can be its puncture and catheterization, or damage to the vessel wall is the first cause of the triad. Hypercoagulation, the second cause of the triad, occurs in elderly patients with high body mass index, cancer, and thrombophilia, and increases the risk of thrombosis.^7,8^ Scientific data demonstrating the impact of the third cause of the triad - venous stasis on the risk of thrombosis - is not enough.^2^ At the same time, it is known that a number of medical interventions, such as intra- and postoperative mechanical ventilation, and patient position, can increase the risk of IJV thrombosis after its catheterization.^2,9^ In particular, cases of catheterized IJV thrombosis occurred in mechanically ventilated patients 8 times more often than those who did not require a ventilator.^9^ A possible reason for this fact may be the effect of mechanical ventilation-induced continuous positive airway pressure on blood flow velocity in the IJV.

In this regard, the purpose of our study was to evaluate the dynamics of changes in the blood flow velocity in the IJV under conditions of changing intrapulmonary pressure and gravitational influence in healthy volunteers.

## Methods

Color duplex ultrasonography of the right IJV was performed in 25 healthy participants of both genders. Written informed consent to participate in this research was obtained from all participants. Eligibility criteria in study was: 1) age 18 years and older, 2) absence of diseases, 3) absence of signs of hypovolemia (tachycardia, hypotension, decrease in the rate of diuresis, 4) absence of medication, 5) absence of alcohol intake for at least 5 days, 6) presence of an informed consent to participate in this study. Withdrawal criteria: 1) non-compliance with the inclusion criteria, 2) obesity.

Ultrasound scanning was performed with portable ultrasound scanner (Mindray, TE7, China) with phased array transducer P4-2s (1.3 – 4.7 MHz). The dopplergram of the blood flow spectrum was recorded in the right IJV in the horizontal position of the volunteers, and also 3 minutes after after the volunteers had attained the new position due to the lifting the bed back section at the 30°. We recorded the maximum (Max) and minimum (Min) linear of blood flow velocity (cm/s). In each position, the measurements were carried out sequentially at positive end-expiratory pressure (PEEP) values of 0, 5, 10 cmH2O after 3 minutes by process of changing the level of PEEP. To create positive breathing in the respiratory tract, a face mask was used, which was connected to a ventilator via a breathing circuit (Mindray CV300, China): ventilator mode – constant positive airway pressure (CPAP).

Statistical analysis was carried out using the IBM SPSS Statistics. Quantitative data were presented as arithmetic mean (M), standard deviation (SD) and compared using the Wilcoxon rank-sum test. The threshold for statistical significance was set at p<0.05. Quantitative indicators were assessed for compliance with the normal distribution using the Shapiro-Wilk test (when the number of subjects was less than 50). In the absence of a normal distribution, quantitative data were described using the median (Me) and the lower and upper quartiles (Q1 – Q3).

## Results

The research was conducted in 25 healthy volunteers (60% male and 40% female). Their characteristics are presented in Table 1.

**Table 1.** The characteristics of healthy participants.

| Indices | Me | Q1-Q3 | Min | Max |
| --- | --- | --- | --- | --- |
| Age, y | 24 | 23-25 | 18 | 35 |
| Body height,m | 1.72 | 1.66-1.76 | 1.50 | 1.90 |
| Weight, kg | 67.50 | 55.50-75.00 | 44.00 | 115.00 |
| Body mass index, kg/m<sup>2</sup> | 22.52 | 19.93-25.54 | 17.30 | 32.19 |

In each subject, the blood flow in the internal jugular vein was estimated under conditions of a changing intrapulmonary pressure and gravitational effect (Fig. 1-3).

**Figure 1.**
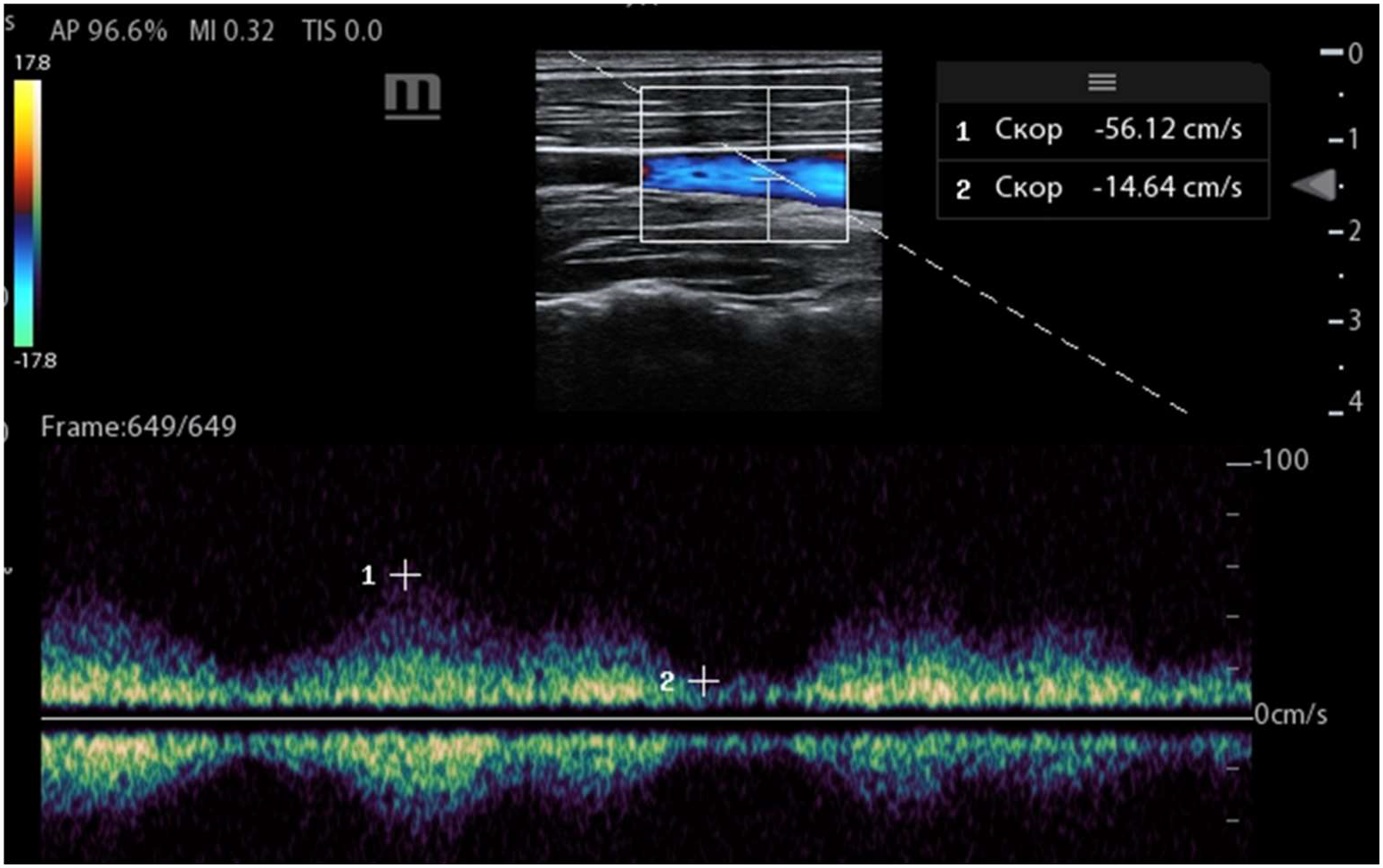
Right internal jugular venous-flow velocity of volunteer in the supine position: PEEP=0 cmH2O.

**Figure 2.**
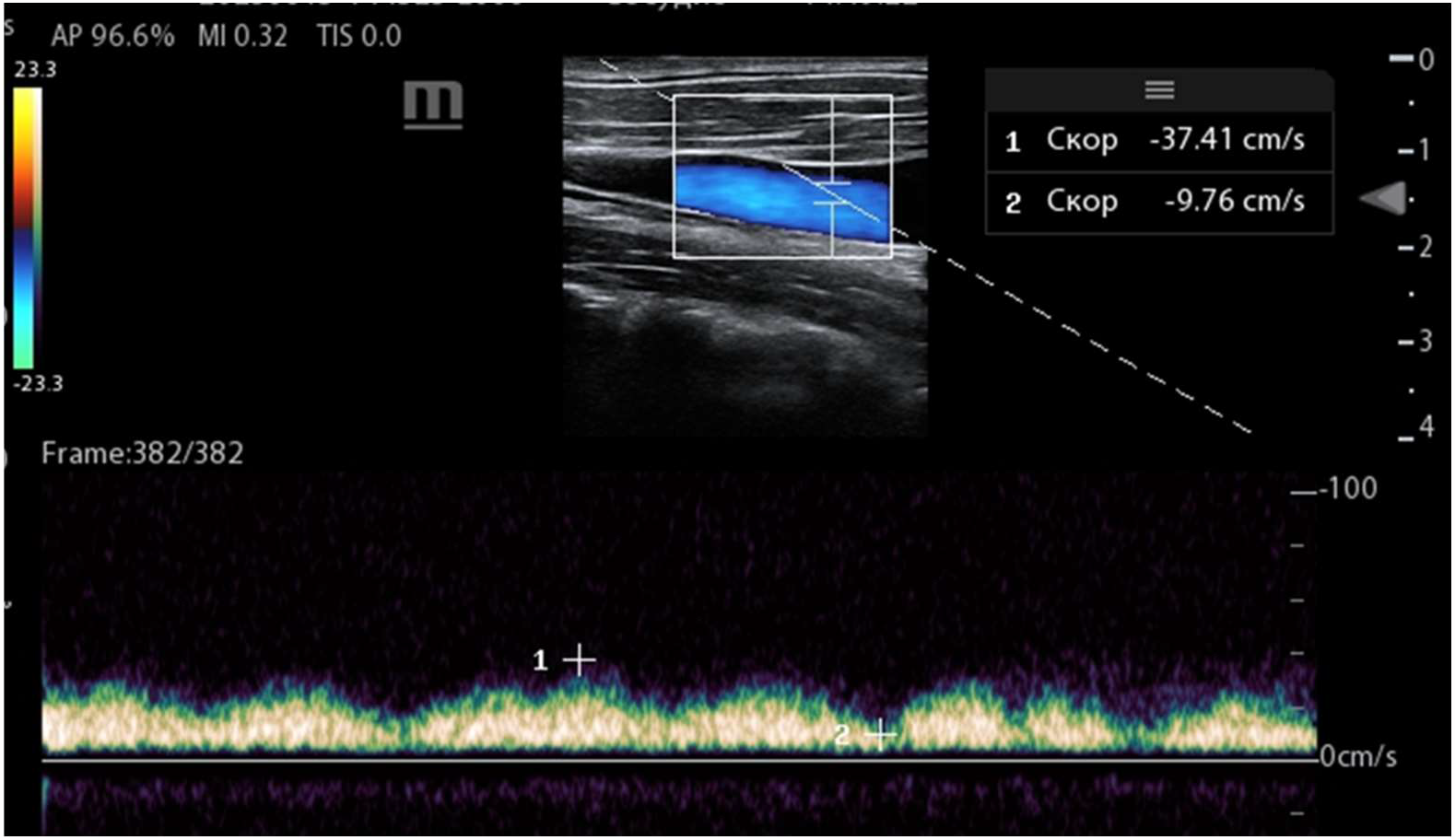
Right internal jugular venous-flow velocity of volunteer in the supine position: PEEP=10 cmH2O.

**Figure 3.**
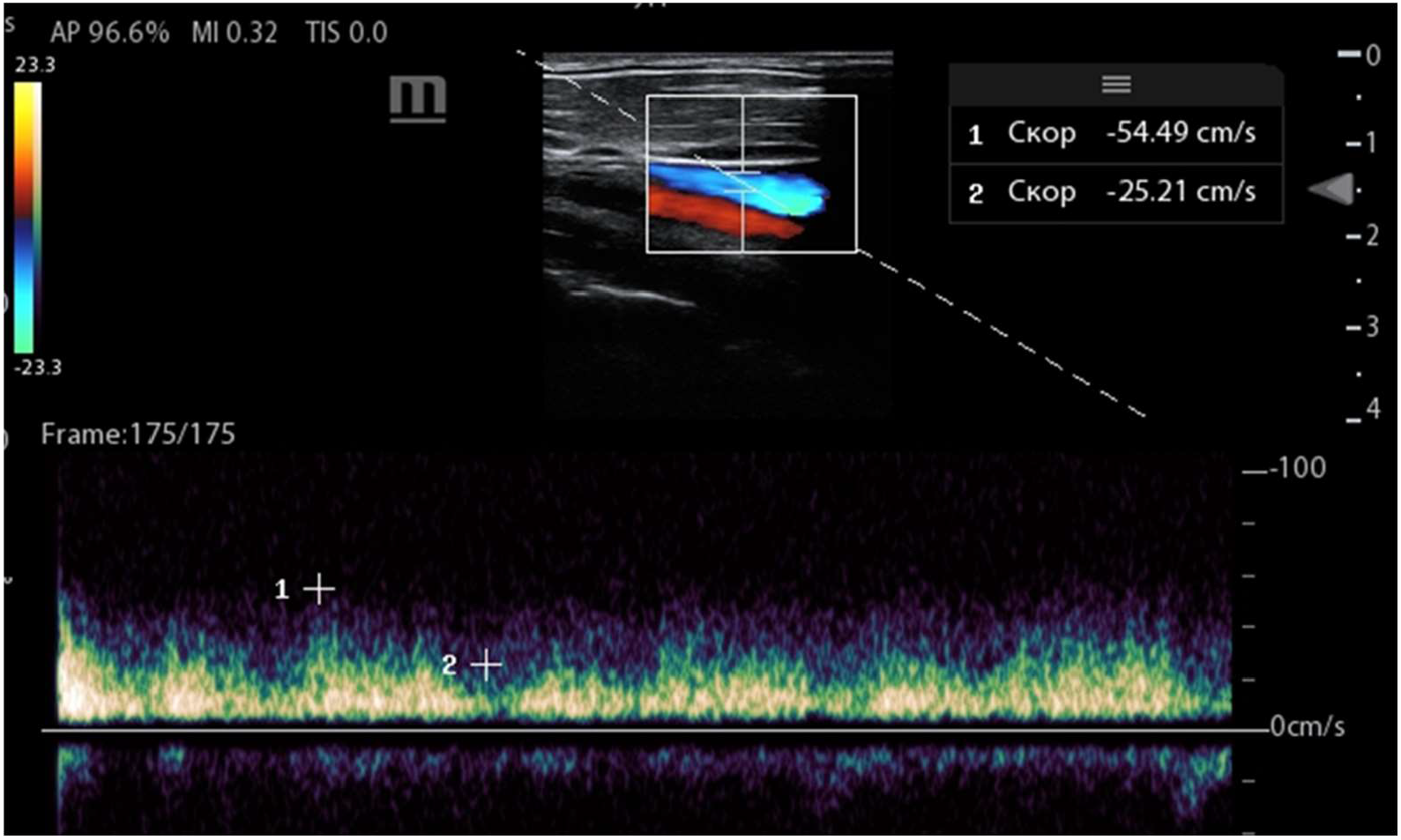
Right internal jugular venous-flow velocity of volunteer after head elevation at 30°: PEEP=10 cmH2O.

The results of the measurement of the IJV blood flow obtained from volunteers in a horizontal supine position under conditions of changing intrapulmonary pressure are shown in Table 2.

**Table 2.**
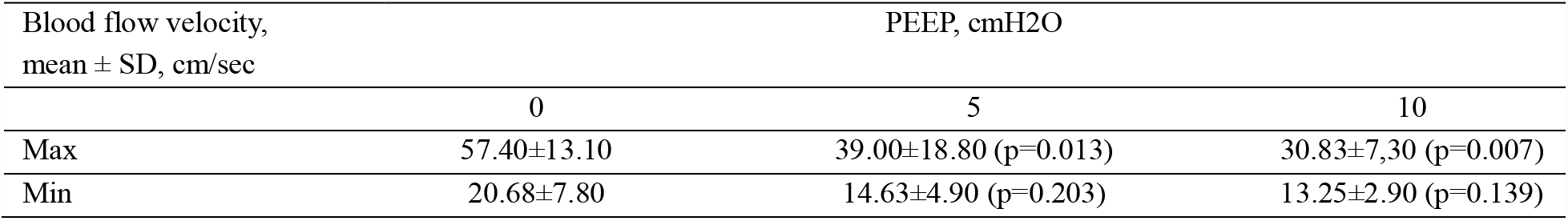
The values of blood flow velocity in the right IJV under conditions of different PEEP

Submitted data is demonstrating a decrease in blood flow velocity in PEEP 5 and 10 cmH2O comparatively base-line values (PEEP 0). At the same time, statistically significant decrease is evaluated only for maximum velocity values. The influence 30° head elevation on IJV blood flow velocity in volunteers with different levels of PEEP are shown in Table 3.

**Table 3.** The values of blood flow velocity in the right IJV under conditions of different PEEP at 30° head elevation

| Blood flow velocity,<br>mean ± SD, cm/sec | PEEP, cmH<sub>2</sub>O | | | |
| --- | --- | --- | --- | --- |
|  | 5 | p-value | 10 | p-value |
| Max | 50.09±23.30 | 0.017 | 42.24±14,4 | 0.047 |
| Min | 18.17±6.30 | 0.008 | 16.24±5.2 | 0.086 |

The blood flow velocity values are increased after 30° head elevation in PEEP 5 and 10 cmH2O in comparison to 0° values. Increasing blood flow in PEEP 5 was statistically significant for maximum and minimum velocity and in PEEP 10 cmH2O — only for maximum velocity.

## Discussion

Obtained data demonstrated that negatively related influence of continuous positive airway pressure on blood flow in internal jugular vein. Mean values of blood flow velocity are decreased with pressure increment in airways. In that way PEEP increase from 0 to 10 cmH2O is go with maximal venous-flow velocity decrease on 46%, and minimal – on 34%. This influence can be important for a patient, who have continued mechanical ventilation and come to be a potential reason of multiplication risk thrombosis IJV, along with factors, like infection and coagulopathy.^9^ However, at the present day is not exact understanding what blood flow decrease in internal jugular vein and what time window is critical for pathogenic mechanism. Only single clinical cases have been demonstrated, that decrease of blood flow in central veins can maximize thrombotic risk. For example, Kimito Minami et al. gave a demonstration of thrombus formation in catheterized IJV after two hours of being patient in defense attitude, which leads to blood flow rate depression in vein to 8 cm/sec.^2^ James Pavela et al. have been reported about decrease of peak blood flow velocity in astronauts, who has been presented in longtime missing of gravitational effect in long duration space mission, which leads to increased echogenicity in the lumen of an intact vein.^1^ He has been made a prediction that in the presence of other factors, decrease of blood flow rate in IJV can be able to increase thrombus risk.

A big scientific data may be helpful for impact assessment of a positive airway pressure on the blood flow velocity changes in the IJV, and also for critical values increasing the risk of thrombosis. A well-known maneuver – increasing head elevation angle – has been already used for increased venous outflow from the cranial cavity^10^ and the dural venous pressure assessment^11^. We were the first to use this maneuver to estimate blood flow velocity in the IJV in people with continuous positive pressure in thorax. Our research has demonstrated, that controlling gravitational effect by means of increasing head of a bed angle allows to reduce PEEP influence. These results suggest that this maneuver can be potentially helpful in clinical practice. At the same time, extension studies help wanted for approving this suggestion. Continuing work will focus on sensitivity analysis of blood flow velocity in patients with catheterizing IJV in mechanical ventilation on thrombotic risk.

The constant positive airway pressure 5 and more cmH2O in a person in the horizontal supine position, decreasing blood flow velocity in IJV. The 30° head elevation eliminates this effect.

## Sources of Funding

This research received no external funding

## Disclosures

None

## Notes

### Competing Interest Statement

The authors have declared no competing interest.

